## Supplementary Figures S1-S3 for "A Systematic Review of the Distribution and Prevalence of Viruses Detected in the *Peromyscus maniculatus* Species Complex (Rodentia: Cricetidae)"

**Figure S1 (next page):** Phylogeny of the hantavirus S genome segment based on a maximum-likelihood alignment of hantavirus nucleotide sequences collected from *Peromyscus maniculatus*. The sequence label in red is the Sin Nombre reference sequence from NCBI. *P. maniculatus* sequences highlighted in blue are used for orientation across the S, M, and L segments, since these sequences are all derived from a single study (1).

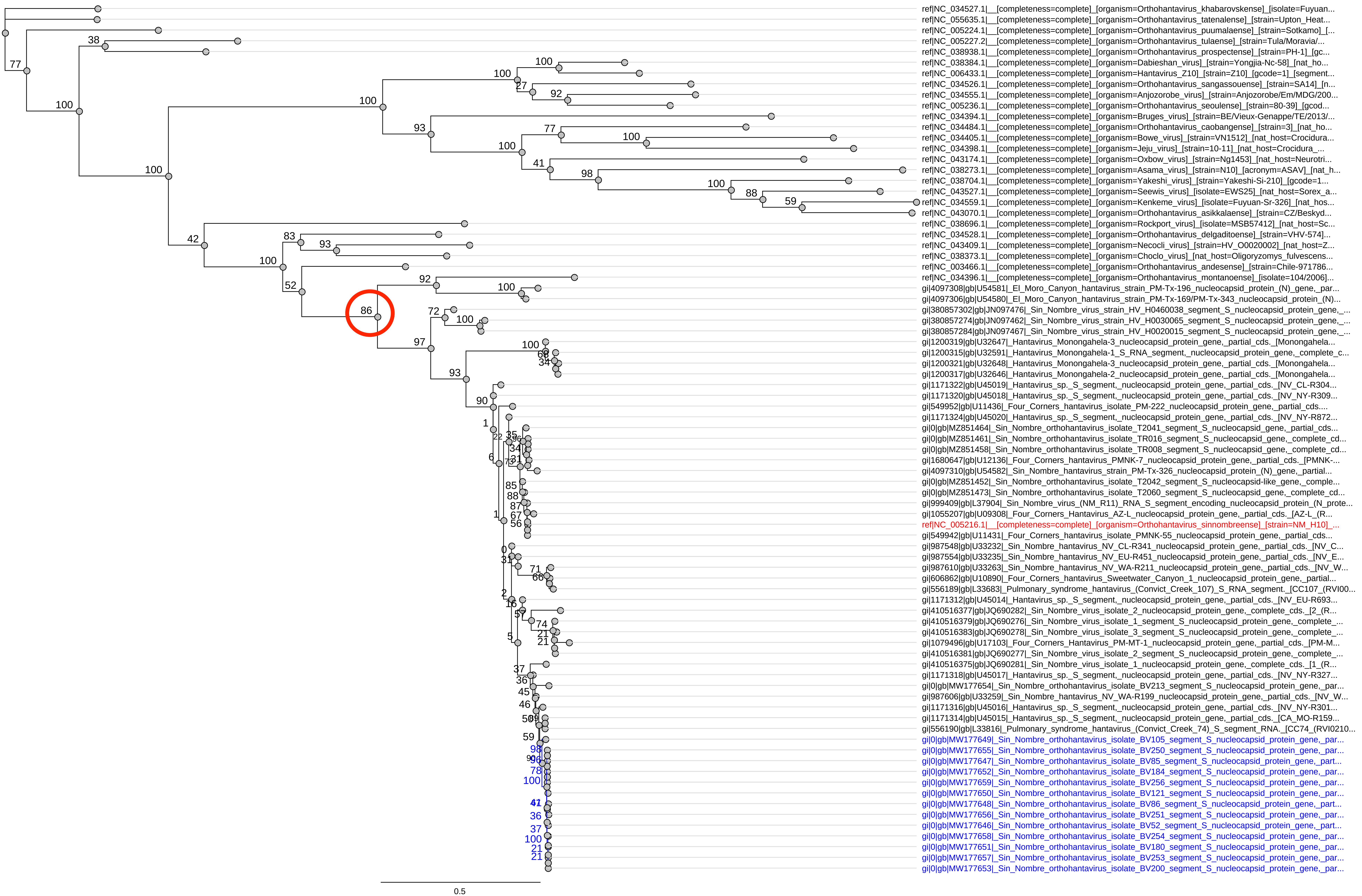
ref|NC_034527.1|__[completeness=complete]_[organism=Orthohantavirus_khabarovskense]_[isolate=Fuyuan...
ref|NC_055635.1|__[completeness=complete]_[organism=Orthohantavirus_tatenalense]_[strain=Upton_Heat...
ref|NC_005224.1|__[completeness=complete]_[organism=Orthohantavirus_puumalaense]_[strain=Sotkamo]_[...
ref|NC_005227.2|__[completeness=complete]_[organism=Orthohantavirus_tulaense]_[strain=Tula/Moravia/...
ref|NC_038938.1|__[completeness=complete]_[organism=Orthohantavirus_prospectense]_[strain=PH-1]_[gc...
ref|NC_038384.1|__[completeness=complete]_[organism=Dabieshan_virus]_[strain=Yongjia-Nc-58]_[nat_ho...
ref|NC_006433.1|__[completeness=complete]_[organism=Hantavirus_Z10]_[strain=Z10]_[gcode=1]_[segment...
ref|NC_034526.1|__[completeness=complete]_[organism=Orthohantavirus_sangassouense]_[strain=SA14]_[n...
ref|NC_034555.1|__[completeness=complete]_[organism=Anjozorobe_virus]_[strain=Anjozorobe/Em/MDG/200...
ref|NC_005236.1|__[completeness=complete]_[organism=Orthohantavirus_seoulense]_[strain=80-39]_[gcod...
ref|NC_034394.1|__[completeness=complete]_[organism=Bruges_virus]_[strain=BE/Vieux-Genappe/TE/2013/...
ref|NC_034484.1|__[completeness=complete]_[organism=Orthohantavirus_caobangense]_[strain=3]_[nat_ho...
ref|NC_034405.1|__[completeness=complete]_[organism=Bowe_virus]_[strain=VN1512]_[nat_host=Crocidura...
ref|NC_034398.1|__[completeness=complete]_[organism=Jeju_virus]_[strain=10-11]_[nat_host=Crocidura_...
ref|NC_043174.1|__[completeness=complete]_[organism=Oxbow_virus]_[strain=Ng1453]_[nat_host=Neurotri...
ref|NC_038273.1|__[completeness=complete]_[organism=Asama_virus]_[strain=N10]_[acronym=ASAV]_[nat_h...
ref|NC_038704.1|__[completeness=complete]_[organism=Yakeshi_virus]_[strain=Yakeshi-Si-210]_[gcode=1...
ref|NC_043527.1|__[completeness=complete]_[organism=Seewis_virus]_[isolate=EWS25]_[nat_host=Sorex_a...
ref|NC_034559.1|__[completeness=complete]_[organism=Kenkeme_virus]_[isolate=Fuyuan-Sr-326]_[nat_hos...
ref|NC_043070.1|__[completeness=complete]_[organism=Orthohantavirus_asikkalaense]_[strain=CZ/Beskyd...
ref|NC_038696.1|__[completeness=complete]_[organism=Rockport_virus]_[isolate=MSB57412]_[nat_host=Sc...
ref|NC_034528.1|__[completeness=complete]_[organism=Orthohantavirus_delgaditoense]_[strain=VHV-574]...
ref|NC_043409.1|__[completeness=complete]_[organism=Necocli_virus]_[strain=HV_O0020002]_[nat_host=Z...
ref|NC_038373.1|__[completeness=complete]_[organism=Choclo_virus]_[nat_host=Oligoryzomys_fulvescens...
ref|NC_003466.1|__[completeness=complete]_[organism=Orthohantavirus_andesense]_[strain=Chile-971786...
ref|NC_034396.1|__[completeness=complete]_[organism=Orthohantavirus_montanoense]_[isolate=104/2006]...
gi|4097308|gb|U54581|_El_Moro_Canyon_hantavirus_strain_PM-Tx-196_nucleocapsid_protein_(N)_gene,_par...
gi|4097306|gb|U54580|_El_Moro_Canyon_hantavirus_strain_PM-Tx-169/PM-Tx-343_nucleocapsid_protein_(N)...
gi|380857302|gb|JN097476|_Sin_Nombre_virus_strain_HV_H0460038_segment_S_nucleocapsid_protein_gene,_...
gi|380857274|gb|JN097462|_Sin_Nombre_virus_strain_HV_H0030065_segment_S_nucleocapsid_protein_gene,_...
gi|380857284|gb|JN097467|_Sin_Nombre_virus_strain_HV_H0020015_segment_S_nucleocapsid_protein_gene,_...
gi|1200319|gb|U32647|_Hantavirus_Monongahela-3_nucleocapsid_protein_gene,_partial_cds._[Monongahela...
gi|1200315|gb|U32591|_Hantavirus_Monongahela-1_S_RNA_segment,_nucleocapsid_protein_gene,_complete_c...
gi|1200321|gb|U32648|_Hantavirus_Monongahela-3_nucleocapsid_protein_gene,_partial_cds._[Monongahela...
gi|1200317|gb|U32646|_Hantavirus_Monongahela-2_nucleocapsid_protein_gene,_partial_cds._[Monongahela...
gi|1171322|gb|U45019|_Hantavirus_sp._S_segment,_nucleocapsid_protein_gene,_partial_cds._[NV_CL-R304...
gi|1171320|gb|U45018|_Hantavirus_sp._S_segment,_nucleocapsid_protein_gene,_partial_cds._[NV_NY-R309...
gi|549952|gb|U11436|_Four_Corners_hantavirus_isolate_PM-222_nucleocapsid_protein_gene,_partial_cds....
gi|1171324|gb|U45020|_Hantavirus_sp._S_segment,_nucleocapsid_protein_gene,_partial_cds._[NV_NY-R872...
gi|0|gb|MZ851464|_Sin_Nombre_orthohantavirus_isolate_T2041_segment_S_nucleocapsid_gene,_partial_cds...
gi|0|gb|MZ851461|_Sin_Nombre_orthohantavirus_isolate_TR016_segment_S_nucleocapsid_gene,_complete_cd...
gi|0|gb|MZ851458|_Sin_Nombre_orthohantavirus_isolate_TR008_segment_S_nucleocapsid_gene,_complete_cd...
gi|1680647|gb|U12136|_Four_Corners_hantavirus_PMNK-7_nucleocapsid_protein_gene,_partial_cds._[PMNK-...
gi|4097310|gb|U54582|_Sin_Nombre_hantavirus_strain_PM-Tx-326_nucleocapsid_protein_(N)_gene,_partial...
gi|0|gb|MZ851452|_Sin_Nombre_orthohantavirus_isolate_T2042_segment_S_nucleocapsid-like_gene,_comple...
gi|0|gb|MZ851473|_Sin_Nombre_orthohantavirus_isolate_T2060_segment_S_nucleocapsid_gene,_complete_cd...
gi|999409|gb|L37904|_Sin_Nombre_virus_(NM_R11)_RNA_S_segment_encoding_nucleocapsid_protein_(N_prote...
gi|1055207|gb|U09308|_Four_Corners_Hantavirus_AZ-L_nucleocapsid_protein_gene,_partial_cds._[AZ-L_(R...
ref|NC_005216.1|__[completeness=complete]_[organism=Orthohantavirus_sinnombreense]_[strain=NM_H10]_...
gi|549942|gb|U11431|_Four_Corners_hantavirus_isolate_PMNK-55_nucleocapsid_protein_gene,_partial_cds...
gi|987548|gb|U33232|_Sin_Nombre_hantavirus_NV_CL-R341_nucleocapsid_protein_gene,_partial_cds._[NV_C...
gi|987554|gb|U33235|_Sin_Nombre_hantavirus_NV_EU-R451_nucleocapsid_protein_gene,_partial_cds._[NV_E...
gi|987610|gb|U33263|_Sin_Nombre_hantavirus_NV_WA-R211_nucleocapsid_protein_gene,_partial_cds._[NV_W...
gi|606862|gb|U10890|_Four_Corners_hantavirus_Sweetwater_Canyon_1_nucleocapsid_protein_gene,_partial...
gi|556189|gb|L33683|_Pulmonary_syndrome_hantavirus_(Convict_Creek_107)_S_RNA_segment._[CC107_(RVI00...
gi|1171312|gb|U45014|_Hantavirus_sp._S_segment,_nucleocapsid_protein_gene,_partial_cds._[NV_EU-R693...
gi|410516377|gb|JQ690282|_Sin_Nombre_virus_isolate_2_nucleocapsid_protein_gene,_complete_cds._[2_(R...
gi|410516379|gb|JQ690276|_Sin_Nombre_virus_isolate_1_segment_S_nucleocapsid_protein_gene,_complete_...
gi|410516383|gb|JQ690278|_Sin_Nombre_virus_isolate_3_segment_S_nucleocapsid_protein_gene,_complete_...
gi|1079496|gb|U17103|_Four_Corners_Hantavirus_PM-MT-1_nucleocapsid_protein_gene,_partial_cds._[PM-M...
gi|410516381|gb|JQ690277|_Sin_Nombre_virus_isolate_2_segment_S_nucleocapsid_protein_gene,_complete_...
gi|410516375|gb|JQ690281|_Sin_Nombre_virus_isolate_1_nucleocapsid_protein_gene,_complete_cds._[1_(R...
gi|1171318|gb|U45017|_Hantavirus_sp._S_segment,_nucleocapsid_protein_gene,_partial_cds._[NV_NY-R327...
gi|0|gb|MW177654|_Sin_Nombre_orthohantavirus_isolate_BV213_segment_S_nucleocapsid_protein_gene,_par...
gi|987606|gb|U33259|_Sin_Nombre_hantavirus_NV_WA-R199_nucleocapsid_protein_gene,_partial_cds._[NV_W...
gi|1171316|gb|U45016|_Hantavirus_sp._S_segment,_nucleocapsid_protein_gene,_partial_cds._[NV_NY-R301...
gi|1171314|gb|U45015|_Hantavirus_sp._S_segment,_nucleocapsid_protein_gene,_partial_cds._[CA_MO-R159...
gi|556190|gb|L33816|_Pulmonary_syndrome_hantavirus_(Convict_Creek_74)_S_segment_RNA._[CC74_(RVI0210...
gi|0|gb|MW177649|_Sin_Nombre_orthohantavirus_isolate_BV105_segment_S_nucleocapsid_protein_gene,_par...
gi|0|gb|MW177655|_Sin_Nombre_orthohantavirus_isolate_BV250_segment_S_nucleocapsid_protein_gene,_par...
gi|0|gb|MW177647|_Sin_Nombre_orthohantavirus_isolate_BV85_segment_S_nucleocapsid_protein_gene,_part...
gi|0|gb|MW177652|_Sin_Nombre_orthohantavirus_isolate_BV184_segment_S_nucleocapsid_protein_gene,_par...
gi|0|gb|MW177659|_Sin_Nombre_orthohantavirus_isolate_BV256_segment_S_nucleocapsid_protein_gene,_par...
gi|0|gb|MW177650|_Sin_Nombre_orthohantavirus_isolate_BV121_segment_S_nucleocapsid_protein_gene,_par...
gi|0|gb|MW177648|_Sin_Nombre_orthohantavirus_isolate_BV86_segment_S_nucleocapsid_protein_gene,_part...
gi|0|gb|MW177656|_Sin_Nombre_orthohantavirus_isolate_BV251_segment_S_nucleocapsid_protein_gene,_par...
gi|0|gb|MW177646|_Sin_Nombre_orthohantavirus_isolate_BV52_segment_S_nucleocapsid_protein_gene,_part...
gi|0|gb|MW177658|_Sin_Nombre_orthohantavirus_isolate_BV254_segment_S_nucleocapsid_protein_gene,_par...
gi|0|gb|MW177651|_Sin_Nombre_orthohantavirus_isolate_BV180_segment_S_nucleocapsid_protein_gene,_par...
gi|0|gb|MW177657|_Sin_Nombre_orthohantavirus_isolate_BV253_segment_S_nucleocapsid_protein_gene,_par...
gi|0|gb|MW177653|_Sin_Nombre_orthohantavirus_isolate_BV200_segment_S_nucleocapsid_protein_gene,_par...
0.5

**Figure S2 (next page):** Phylogeny of the hantavirus M genome segment based on a maximum-likelihood alignment of hantavirus nucleotide sequences collected from *Peromyscus maniculatus*. The sequence label in red is the Sin Nombre reference sequence from NCBI. *P. maniculatus* sequences highlighted in blue are used for orientation across the S, M, and L segments, since these sequences are all derived from a single study (1).

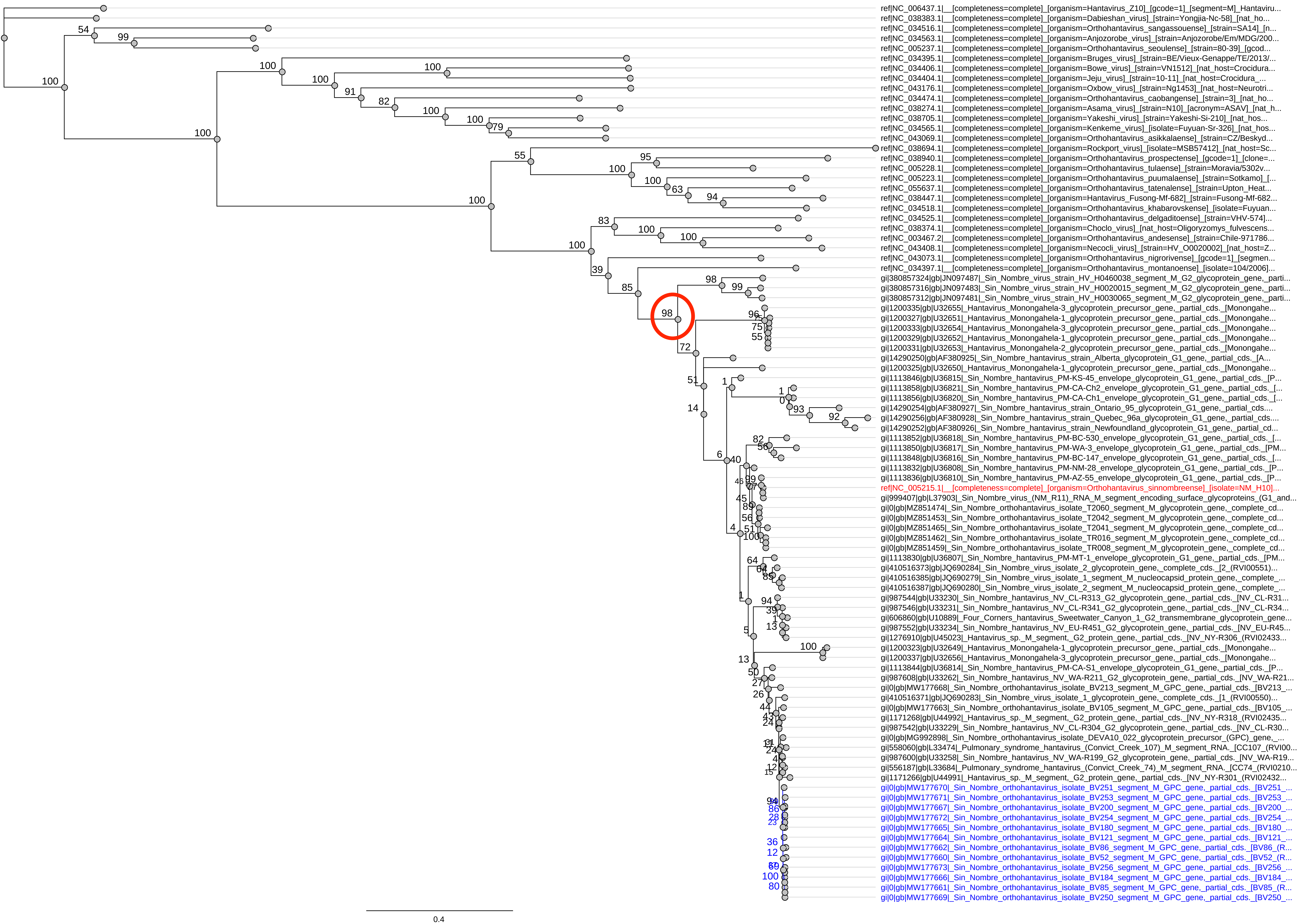

ref|NC_006437.1|__[completeness=complete]_[organism=Hantavirus_Z10]_[gcode=1]_[segment=M]_Hantaviru...
ref|NC_038383.1|__[completeness=complete]_[organism=Dabieshan_virus]_[strain=Yongjia-Nc-58]_[nat_ho...
ref|NC_034516.1|__[completeness=complete]_[organism=Orthohantavirus_sangassouense]_[strain=SA14]_[n...
ref|NC_034563.1|__[completeness=complete]_[organism=Anjozorobe_virus]_[strain=Anjozorobe/Em/MDG/200...
ref|NC_005237.1|__[completeness=complete]_[organism=Orthohantavirus_seoulense]_[strain=80-39]_[gcod...
ref|NC_034395.1|__[completeness=complete]_[organism=Bruges_virus]_[strain=BE/Vieux-Genappe/TE/2013/...
ref|NC_034406.1|__[completeness=complete]_[organism=Bowe_virus]_[strain=VN1512]_[nat_host=Crocidura...
ref|NC_034404.1|__[completeness=complete]_[organism=Jeju_virus]_[strain=10-11]_[nat_host=Crocidura_...
ref|NC_043176.1|__[completeness=complete]_[organism=Oxbow_virus]_[strain=Ng1453]_[nat_host=Neurotri...
ref|NC_034474.1|__[completeness=complete]_[organism=Orthohantavirus_caobangense]_[strain=3]_[nat_ho...
ref|NC_038274.1|__[completeness=complete]_[organism=Asama_virus]_[strain=N10]_[acronym=ASAV]_[nat_h...
ref|NC_038705.1|__[completeness=complete]_[organism=Yakeshi_virus]_[strain=Yakeshi-Si-210]_[nat_hos...
ref|NC_034565.1|__[completeness=complete]_[organism=Kenkeme_virus]_[isolate=Fuyuan-Sr-326]_[nat_hos...
ref|NC_043069.1|__[completeness=complete]_[organism=Orthohantavirus_asikkalaense]_[strain=CZ/Beskyd...
ref|NC_038694.1|__[completeness=complete]_[organism=Rockport_virus]_[isolate=MSB57412]_[nat_host=Sc...
ref|NC_038940.1|__[completeness=complete]_[organism=Orthohantavirus_prospectense]_[gcode=1]_[clone=...
ref|NC_005228.1|__[completeness=complete]_[organism=Orthohantavirus_tulaense]_[strain=Moravia/5302v...
ref|NC_005223.1|__[completeness=complete]_[organism=Orthohantavirus_puumalaense]_[strain=Sotkamo]_[...
ref|NC_055637.1|__[completeness=complete]_[organism=Orthohantavirus_tatenalense]_[strain=Upton_Heat...
ref|NC_038447.1|__[completeness=complete]_[organism=Hantavirus_Fusong-Mf-682]_[strain=Fusong-Mf-682...
ref|NC_034518.1|__[completeness=complete]_[organism=Orthohantavirus_khabarovskense]_[isolate=Fuyuan...
ref|NC_034525.1|__[completeness=complete]_[organism=Orthohantavirus_delgaditoense]_[strain=VHV-574]...
ref|NC_038374.1|__[completeness=complete]_[organism=Choclo_virus]_[nat_host=Oligoryzomys_fulvescens...
ref|NC_003467.2|__[completeness=complete]_[organism=Orthohantavirus_andesense]_[strain=Chile-971786...
ref|NC_043408.1|__[completeness=complete]_[organism=Necocli_virus]_[strain=HV_O0020002]_[nat_host=Z...
ref|NC_043073.1|__[completeness=complete]_[organism=Orthohantavirus_nigrorivense]_[gcode=1]_[segmen...
ref|NC_034397.1|__[completeness=complete]_[organism=Orthohantavirus_montanoense]_[isolate=104/2006]...
gi|380857324|gb|JN097487|_Sin_Nombre_virus_strain_HV_H0460038_segment_M_G2_glycoprotein_gene,_parti...
gi|380857316|gb|JN097483|_Sin_Nombre_virus_strain_HV_H0020015_segment_M_G2_glycoprotein_gene,_parti...
gi|380857312|gb|JN097481|_Sin_Nombre_virus_strain_HV_H0030065_segment_M_G2_glycoprotein_gene,_parti...
gi|1200335|gb|U32655|_Hantavirus_Monongahela-3_glycoprotein_precursor_gene,_partial_cds._[Monongahe...
gi|1200327|gb|U32651|_Hantavirus_Monongahela-1_glycoprotein_precursor_gene,_partial_cds._[Monongahe...
gi|1200333|gb|U32654|_Hantavirus_Monongahela-3_glycoprotein_precursor_gene,_partial_cds._[Monongahe...
gi|1200329|gb|U32652|_Hantavirus_Monongahela-1_glycoprotein_precursor_gene,_partial_cds._[Monongahe...
gi|1200331|gb|U32653|_Hantavirus_Monongahela-2_glycoprotein_precursor_gene,_partial_cds._[Monongahe...
gi|14290250|gb|AF380925|_Sin_Nombre_hantavirus_strain_Alberta_glycoprotein_G1_gene,_partial_cds._[A...
gi|1200325|gb|U32650|_Hantavirus_Monongahela-1_glycoprotein_precursor_gene,_partial_cds._[Monongahe...
gi|1113846|gb|U36815|_Sin_Nombre_hantavirus_PM-KS-45_envelope_glycoprotein_G1_gene,_partial_cds._[P...
gi|1113858|gb|U36821|_Sin_Nombre_hantavirus_PM-CA-Ch2_envelope_glycoprotein_G1_gene,_partial_cds._[...
gi|1113856|gb|U36820|_Sin_Nombre_hantavirus_PM-CA-Ch1_envelope_glycoprotein_G1_gene,_partial_cds._[...
gi|14290254|gb|AF380927|_Sin_Nombre_hantavirus_strain_Ontario_95_glycoprotein_G1_gene,_partial_cds....
gi|14290256|gb|AF380928|_Sin_Nombre_hantavirus_strain_Quebec_96a_glycoprotein_G1_gene,_partial_cds....
gi|14290252|gb|AF380926|_Sin_Nombre_hantavirus_strain_Newfoundland_glycoprotein_G1_gene,_partial_cd...
gi|1113852|gb|U36818|_Sin_Nombre_hantavirus_PM-BC-530_envelope_glycoprotein_G1_gene,_partial_cds._[...
gi|1113850|gb|U36817|_Sin_Nombre_hantavirus_PM-WA-3_envelope_glycoprotein_G1_gene,_partial_cds._[PM...
gi|1113848|gb|U36816|_Sin_Nombre_hantavirus_PM-BC-147_envelope_glycoprotein_G1_gene,_partial_cds._[...
gi|1113832|gb|U36808|_Sin_Nombre_hantavirus_PM-NM-28_envelope_glycoprotein_G1_gene,_partial_cds._[P...
gi|1113836|gb|U36810|_Sin_Nombre_hantavirus_PM-AZ-55_envelope_glycoprotein_G1_gene,_partial_cds._[P...
ref|NC_005215.1|__[completeness=complete]_[organism=Orthohantavirus_sinnombreense]_[isolate=NM_H10]...
gi|999407|gb|L37903|_Sin_Nombre_virus_(NM_R11)_RNA_M_segment_encoding_surface_glycoproteins_(G1_and...
gi|0|gb|MZ851474|_Sin_Nombre_orthohantavirus_isolate_T2060_segment_M_glycoprotein_gene,_complete_cd...
gi|0|gb|MZ851453|_Sin_Nombre_orthohantavirus_isolate_T2042_segment_M_glycoprotein_gene,_complete_cd...
gi|0|gb|MZ851465|_Sin_Nombre_orthohantavirus_isolate_T2041_segment_M_glycoprotein_gene,_complete_cd...
gi|0|gb|MZ851462|_Sin_Nombre_orthohantavirus_isolate_TR016_segment_M_glycoprotein_gene,_complete_cd...
gi|0|gb|MZ851459|_Sin_Nombre_orthohantavirus_isolate_TR008_segment_M_glycoprotein_gene,_complete_cd...
gi|1113830|gb|U36807|_Sin_Nombre_hantavirus_PM-MT-1_envelope_glycoprotein_G1_gene,_partial_cds._[PM...
gi|410516373|gb|JQ690284|_Sin_Nombre_virus_isolate_2_glycoprotein_gene,_complete_cds._[2_(RVI00551)...
gi|410516385|gb|JQ690279|_Sin_Nombre_virus_isolate_1_segment_M_nucleocapsid_protein_gene,_complete_...
gi|410516387|gb|JQ690280|_Sin_Nombre_virus_isolate_2_segment_M_nucleocapsid_protein_gene,_complete_...
gi|987544|gb|U33230|_Sin_Nombre_hantavirus_NV_CL-R313_G2_glycoprotein_gene,_partial_cds._[NV_CL-R31...
gi|987546|gb|U33231|_Sin_Nombre_hantavirus_NV_CL-R341_G2_glycoprotein_gene,_partial_cds._[NV_CL-R34...
gi|606860|gb|U10889|_Four_Corners_hantavirus_Sweetwater_Canyon_1_G2_transmembrane_glycoprotein_gene...
gi|987552|gb|U33234|_Sin_Nombre_hantavirus_NV_EU-R451_G2_glycoprotein_gene,_partial_cds._[NV_EU-R45...
gi|1276910|gb|U45023|_Hantavirus_sp._M_segment,_G2_protein_gene,_partial_cds._[NV_NY-R306_(RVI02433...
gi|1200323|gb|U32649|_Hantavirus_Monongahela-1_glycoprotein_precursor_gene,_partial_cds._[Monongahe...
gi|1200337|gb|U32656|_Hantavirus_Monongahela-3_glycoprotein_precursor_gene,_partial_cds._[Monongahe...
gi|1113844|gb|U36814|_Sin_Nombre_hantavirus_PM-CA-S1_envelope_glycoprotein_G1_gene,_partial_cds._[P...
gi|987608|gb|U33262|_Sin_Nombre_hantavirus_NV_WA-R211_G2_glycoprotein_gene,_partial_cds._[NV_WA-R21...
gi|0|gb|MW177668|_Sin_Nombre_orthohantavirus_isolate_BV213_segment_M_GPC_gene,_partial_cds._[BV213_...
gi|410516371|gb|JQ690283|_Sin_Nombre_virus_isolate_1_glycoprotein_gene,_complete_cds._[1_(RVI00550)...
gi|0|gb|MW177663|_Sin_Nombre_orthohantavirus_isolate_BV105_segment_M_GPC_gene,_partial_cds._[BV105_...
gi|1171268|gb|U44992|_Hantavirus_sp._M_segment,_G2_protein_gene,_partial_cds._[NV_NY-R318_(RVI02435...
gi|987542|gb|U33229|_Sin_Nombre_hantavirus_NV_CL-R304_G2_glycoprotein_gene,_partial_cds._[NV_CL-R30...
gi|0|gb|MG992898|_Sin_Nombre_orthohantavirus_isolate_DEVA10_022_glycoprotein_precursor_(GPC)_gene,_...
gi|558060|gb|L33474|_Pulmonary_syndrome_hantavirus_(Convict_Creek_107)_M_segment_RNA._[CC107_(RVI00...
gi|987600|gb|U33258|_Sin_Nombre_hantavirus_NV_WA-R199_G2_glycoprotein_gene,_partial_cds._[NV_WA-R19...
gi|556187|gb|L33684|_Pulmonary_syndrome_hantavirus_(Convict_Creek_74)_M_segment_RNA._[CC74_(RVI0210...
gi|1171266|gb|U44991|_Hantavirus_sp._M_segment,_G2_protein_gene,_partial_cds._[NV_NY-R301_(RVI02432...
gi|0|gb|MW177670|_Sin_Nombre_orthohantavirus_isolate_BV251_segment_M_GPC_gene,_partial_cds._[BV251_...
gi|0|gb|MW177671|_Sin_Nombre_orthohantavirus_isolate_BV253_segment_M_GPC_gene,_partial_cds._[BV253_...
gi|0|gb|MW177667|_Sin_Nombre_orthohantavirus_isolate_BV200_segment_M_GPC_gene,_partial_cds._[BV200_...
gi|0|gb|MW177672|_Sin_Nombre_orthohantavirus_isolate_BV254_segment_M_GPC_gene,_partial_cds._[BV254_...
gi|0|gb|MW177665|_Sin_Nombre_orthohantavirus_isolate_BV180_segment_M_GPC_gene,_partial_cds._[BV180_...
gi|0|gb|MW177664|_Sin_Nombre_orthohantavirus_isolate_BV121_segment_M_GPC_gene,_partial_cds._[BV121_...
gi|0|gb|MW177662|_Sin_Nombre_orthohantavirus_isolate_BV86_segment_M_GPC_gene,_partial_cds._[BV86_(R...
gi|0|gb|MW177660|_Sin_Nombre_orthohantavirus_isolate_BV52_segment_M_GPC_gene,_partial_cds._[BV52_(R...
gi|0|gb|MW177673|_Sin_Nombre_orthohantavirus_isolate_BV256_segment_M_GPC_gene,_partial_cds._[BV256_...
gi|0|gb|MW177666|_Sin_Nombre_orthohantavirus_isolate_BV184_segment_M_GPC_gene,_partial_cds._[BV184_...
gi|0|gb|MW177661|_Sin_Nombre_orthohantavirus_isolate_BV85_segment_M_GPC_gene,_partial_cds._[BV85_(R...
gi|0|gb|MW177669|_Sin_Nombre_orthohantavirus_isolate_BV250_segment_M_GPC_gene,_partial_cds._[BV250_...
0.4

**Figure S3 (next page):** Phylogeny of the hantavirus L genome segment based on a maximum-likelihood alignment of hantavirus nucleotide sequences collected from *Peromyscus maniculatus*. The sequence label in red is the Sin Nombre reference sequence from NCBI. *P. maniculatus* sequences highlighted in blue are used for orientation across the S, M, and L segments, since these sequences are all derived from a single study (1).

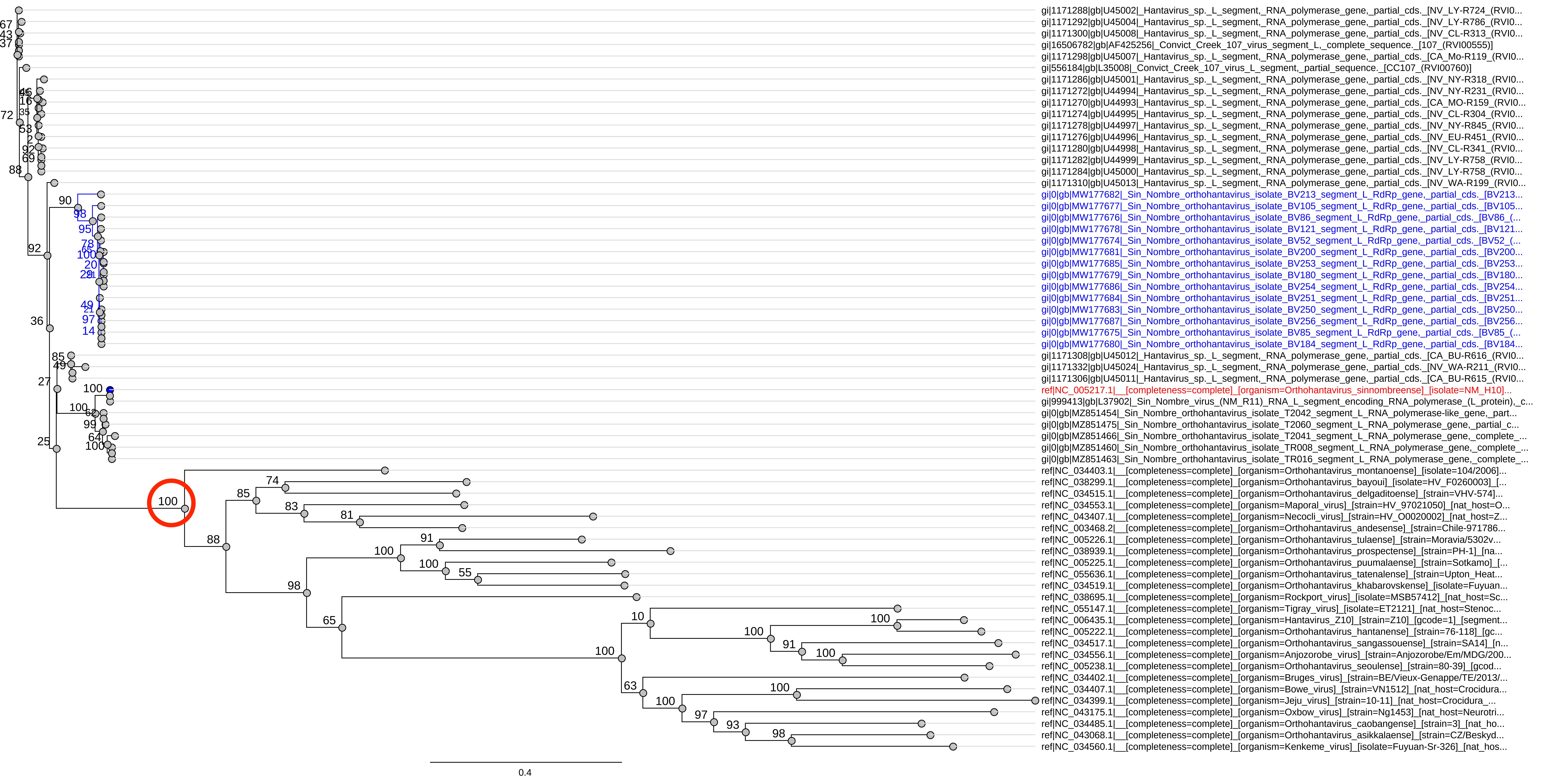
gi|1171288|gb|U45002|_Hantavirus_sp._L_segment,_RNA_polymerase_gene,_partial_cds._[NV_LY-R724_(RVI0...
gi|1171292|gb|U45004|_Hantavirus_sp._L_segment,_RNA_polymerase_gene,_partial_cds._[NV_LY-R786_(RVI0...
gi|1171300|gb|U45008|_Hantavirus_sp._L_segment,_RNA_polymerase_gene,_partial_cds._[NV_CL-R313_(RVI0...
gi|16506782|gb|AF425256|_Convict_Creek_107_virus_segment_L,_complete_sequence._[107_(RVI00555)]
gi|1171298|gb|U45007|_Hantavirus_sp._L_segment,_RNA_polymerase_gene,_partial_cds._[CA_Mo-R119_(RVI0...
gi|556184|gb|L35008|_Convict_Creek_107_virus_L_segment,_partial_sequence._[CC107_(RVI00760)]
gi|1171286|gb|U45001|_Hantavirus_sp._L_segment,_RNA_polymerase_gene,_partial_cds._[NV_NY-R318_(RVI0...
gi|1171272|gb|U44994|_Hantavirus_sp._L_segment,_RNA_polymerase_gene,_partial_cds._[NV_NY-R231_(RVI0...
gi|1171270|gb|U44993|_Hantavirus_sp._L_segment,_RNA_polymerase_gene,_partial_cds._[CA_MO-R159_(RVI0...
gi|1171274|gb|U44995|_Hantavirus_sp._L_segment,_RNA_polymerase_gene,_partial_cds._[NV_CL-R304_(RVI0...
gi|1171278|gb|U44997|_Hantavirus_sp._L_segment,_RNA_polymerase_gene,_partial_cds._[NV_NY-R845_(RVI0...
gi|1171276|gb|U44996|_Hantavirus_sp._L_segment,_RNA_polymerase_gene,_partial_cds._[NV_EU-R451_(RVI0...
gi|1171280|gb|U44998|_Hantavirus_sp._L_segment,_RNA_polymerase_gene,_partial_cds._[NV_CL-R341_(RVI0...
gi|1171282|gb|U44999|_Hantavirus_sp._L_segment,_RNA_polymerase_gene,_partial_cds._[NV_LY-R758_(RVI0...
gi|1171284|gb|U45000|_Hantavirus_sp._L_segment,_RNA_polymerase_gene,_partial_cds._[NV_LY-R758_(RVI0...
gi|1171310|gb|U45013|_Hantavirus_sp._L_segment,_RNA_polymerase_gene,_partial_cds._[NV_WA-R199_(RVI0...
gi|0|gb|MW177682|_Sin_Nombre_orthohantavirus_isolate_BV213_segment_L_RdRp_gene,_partial_cds._[BV213...
gi|0|gb|MW177677|_Sin_Nombre_orthohantavirus_isolate_BV105_segment_L_RdRp_gene,_partial_cds._[BV105...
gi|0|gb|MW177676|_Sin_Nombre_orthohantavirus_isolate_BV86_segment_L_RdRp_gene,_partial_cds._[BV86_(...
gi|0|gb|MW177678|_Sin_Nombre_orthohantavirus_isolate_BV121_segment_L_RdRp_gene,_partial_cds._[BV121...
gi|0|gb|MW177674|_Sin_Nombre_orthohantavirus_isolate_BV52_segment_L_RdRp_gene,_partial_cds._[BV52_(...
gi|0|gb|MW177681|_Sin_Nombre_orthohantavirus_isolate_BV200_segment_L_RdRp_gene,_partial_cds._[BV200...
gi|0|gb|MW177685|_Sin_Nombre_orthohantavirus_isolate_BV253_segment_L_RdRp_gene,_partial_cds._[BV253...
gi|0|gb|MW177679|_Sin_Nombre_orthohantavirus_isolate_BV180_segment_L_RdRp_gene,_partial_cds._[BV180...
gi|0|gb|MW177686|_Sin_Nombre_orthohantavirus_isolate_BV254_segment_L_RdRp_gene,_partial_cds._[BV254...
gi|0|gb|MW177684|_Sin_Nombre_orthohantavirus_isolate_BV251_segment_L_RdRp_gene,_partial_cds._[BV251...
gi|0|gb|MW177683|_Sin_Nombre_orthohantavirus_isolate_BV250_segment_L_RdRp_gene,_partial_cds._[BV250...
gi|0|gb|MW177687|_Sin_Nombre_orthohantavirus_isolate_BV256_segment_L_RdRp_gene,_partial_cds._[BV256...
gi|0|gb|MW177675|_Sin_Nombre_orthohantavirus_isolate_BV85_segment_L_RdRp_gene,_partial_cds._[BV85_(...
gi|0|gb|MW177680|_Sin_Nombre_orthohantavirus_isolate_BV184_segment_L_RdRp_gene,_partial_cds._[BV184...
gi|1171308|gb|U45012|_Hantavirus_sp._L_segment,_RNA_polymerase_gene,_partial_cds._[CA_BU-R616_(RVI0...
gi|1171332|gb|U45024|_Hantavirus_sp._L_segment,_RNA_polymerase_gene,_partial_cds._[NV_WA-R211_(RVI0...
gi|1171306|gb|U45011|_Hantavirus_sp._L_segment,_RNA_polymerase_gene,_partial_cds._[CA_BU-R615_(RVI0...
ref|NC_005217.1|__[completeness=complete]_[organism=Orthohantavirus_sinnombreense]_[isolate=NM_H10]...
gi|999413|gb|L37902|_Sin_Nombre_virus_(NM_R11)_RNA_L_segment_encoding_RNA_polymerase_(L_protein),_c...
gi|0|gb|MZ851454|_Sin_Nombre_orthohantavirus_isolate_T2042_segment_L_RNA_polymerase-like_gene,_part...
gi|0|gb|MZ851475|_Sin_Nombre_orthohantavirus_isolate_T2060_segment_L_RNA_polymerase_gene,_partial_c...
gi|0|gb|MZ851466|_Sin_Nombre_orthohantavirus_isolate_T2041_segment_L_RNA_polymerase_gene,_complete_...
gi|0|gb|MZ851460|_Sin_Nombre_orthohantavirus_isolate_TR008_segment_L_RNA_polymerase_gene,_complete_...
gi|0|gb|MZ851463|_Sin_Nombre_orthohantavirus_isolate_TR016_segment_L_RNA_polymerase_gene,_complete_...
ref|NC_034403.1|__[completeness=complete]_[organism=Orthohantavirus_montanoense]_[isolate=104/2006]...
ref|NC_038299.1|__[completeness=complete]_[organism=Orthohantavirus_bayoui]_[isolate=HV_F0260003]_[...
ref|NC_034515.1|__[completeness=complete]_[organism=Orthohantavirus_delgaditoense]_[strain=VHV-574]...
ref|NC_034553.1|__[completeness=complete]_[organism=Maporal_virus]_[strain=HV_97021050]_[nat_host=O...
ref|NC_043407.1|__[completeness=complete]_[organism=Necocli_virus]_[strain=HV_O0020002]_[nat_host=Z...
ref|NC_003468.2|__[completeness=complete]_[organism=Orthohantavirus_andesense]_[strain=Chile-971786...
ref|NC_005226.1|__[completeness=complete]_[organism=Orthohantavirus_tulaense]_[strain=Moravia/5302v...
ref|NC_038939.1|__[completeness=complete]_[organism=Orthohantavirus_prospectense]_[strain=PH-1]_[na...
ref|NC_005225.1|__[completeness=complete]_[organism=Orthohantavirus_puumalaense]_[strain=Sotkamo]_[...
ref|NC_055636.1|__[completeness=complete]_[organism=Orthohantavirus_tatenalense]_[strain=Upton_Heat...
ref|NC_034519.1|__[completeness=complete]_[organism=Orthohantavirus_khabarovskense]_[isolate=Fuyuan...
ref|NC_038695.1|__[completeness=complete]_[organism=Rockport_virus]_[isolate=MSB57412]_[nat_host=Sc...
ref|NC_055147.1|__[completeness=complete]_[organism=Tigray_virus]_[isolate=ET2121]_[nat_host=Stenoc...
ref|NC_006435.1|__[completeness=complete]_[organism=Hantavirus_Z10]_[strain=Z10]_[gcode=1]_[segment...
ref|NC_005222.1|__[completeness=complete]_[organism=Orthohantavirus_hantanense]_[strain=76-118]_[gc...
ref|NC_034517.1|__[completeness=complete]_[organism=Orthohantavirus_sangassouense]_[strain=SA14]_[n...
ref|NC_034556.1|__[completeness=complete]_[organism=Anjozorobe_virus]_[strain=Anjozorobe/Em/MDG/200...
ref|NC_005238.1|__[completeness=complete]_[organism=Orthohantavirus_seoulense]_[strain=80-39]_[gcod...
ref|NC_034402.1|__[completeness=complete]_[organism=Bruges_virus]_[strain=BE/Vieux-Genappe/TE/2013/...
ref|NC_034407.1|__[completeness=complete]_[organism=Bowe_virus]_[strain=VN1512]_[nat_host=Crocidura...
ref|NC_034399.1|__[completeness=complete]_[organism=Jeju_virus]_[strain=10-11]_[nat_host=Crocidura_...
ref|NC_043175.1|__[completeness=complete]_[organism=Oxbow_virus]_[strain=Ng1453]_[nat_host=Neurotri...
ref|NC_034485.1|__[completeness=complete]_[organism=Orthohantavirus_caobangense]_[strain=3]_[nat_ho...
ref|NC_043068.1|__[completeness=complete]_[organism=Orthohantavirus_asikkalaense]_[strain=CZ/Beskyd...
ref|NC_034560.1|__[completeness=complete]_[organism=Kenkeme_virus]_[isolate=Fuyuan-Sr-326]_[nat_hos...
0.4
